# Fast genotyping of known SNPs through approximate *k*-mer matching

**DOI:** 10.1101/063446

**Authors:** Ariya Shajii, Deniz Yorukoglu, Y. William Yu, Bonnie Berger

## Abstract

**Motivation:** As the volume of next-generation sequencing (NGS) data increases, faster algorithms become necessary. Although speeding up individual components of a sequence analysis pipeline (e.g. read mapping) can reduce the computational cost of analysis, such approaches do not take full advantage of the particulars of a given problem. One problem of great interest, genotyping a known set of variants (e.g. dbSNP or Affymetrix SNPs), is important for characterization of known genetic traits and causative disease variants within an individual, as well as the initial stage of many ancestral and population genomic pipelines (e.g. GWAS).

**Results:** We introduce LAVA (Lightweight Assignment of Variant Alleles), an NGS-based genotyping algorithm for a given set of SNP loci, which takes advantage of the fact that approximate matching of mid-size *k*-mers (with *k* = 32) can typically uniquely identify loci in the human genome without full read alignment. LAVA accurately calls the vast majority of SNPs in dbSNP and Affymetrix’s Genome-Wide Human SNP Array 6.0 up to about an order of magnitude faster than standard NGS genotyping pipelines. For Affymetrix SNPs, LAVA has significantly higher SNP calling accuracy than existing pipelines while using as low as ~5GB of RAM. As such, LAVA represents a scalable computational method for population-level genotyping studies as well as a flexible NGS-based replacement for SNP arrays.

**Availability:** LAVA software is available at http://lava.csail.mit.edu.

**Supplementary information:** Supplementary data are available at *Bioinformatics* online.

## 1 Introduction

One central challenge in genomics is genotyping: given an individual, identifying the locations at which that individual’s genome differs from a reference (Luikart *et al.*, 2003). In this paper, we will primarily focus on SNPs (single nucleotide polymorphisms)—the most frequently used type of human genetic variation in population-level studies (Lancia *et al.*, 2001)—although other potentially-applicable variants include insertions, deletions, short tandem repeats, CNVs (copy number variations), and rearrangements. For simultaneous characterization of large numbers of known SNPs, such as in genome-wide association studies (GWAS), researchers have traditionally turned to allele-specific oligonucleotides (ASO) probes, often adhered onto a DNA microarray to form SNP arrays (Pastinen *et al.*, 2000).

However, there are millions of known SNPs (Sherry *et al.*, 2001), and even the state-of-the-art Affymetrix genome-wide SNP array 6.0 has only 906,000 SNP probes and 946,000 CNV probes. We can instead turn to whole genome sequencing for genotyping. Currently, NGS whole genome sequencing is still relatively more expensive than SNP arrays, but in recent years, sequencing prices have been dropping drastically, going under even the celebrated $1000 mark (Hayden, 2014).

In most NGS-based genotyping pipelines, the first step after sequencing a genome is to map each read to the reference (Li and Durbin, 2009;Langmead and Salzberg, 2012;Yorukoglu *et al.*, 2016). Standard tools for genotyping (e.g. Samtools mpileup (Li *et al.*, 2009) and GATK HaplotypeCaller ( McKenna *et al.*, 2010)) require this mapping information for every read before being able to call variants. Yet despite recent advances in speed (Marco-Sola *et al.*, 2012;Siragusa *et al.*, 2013;Yorukoglu *et al.*, 2016;Zaharia *et al.*, 2011), mapping still remains a computationally expensive step. Furthermore, genotyping pipelines also include variant calling steps, significantly increasing the total runtime.

**Fig. 1.**
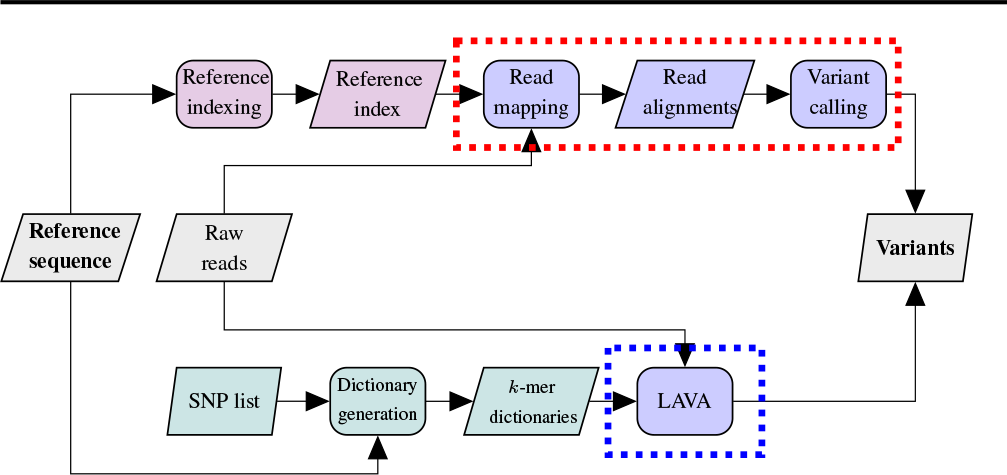
LAVA pipeline (circled in blue) versus conventional genotyping pipeline (circled in red). In contrast to traditional reference indexing (violet), LAVA preprocessing generates *k*-mer dictionaries from a given reference sequence and list of SNPs (teal). Our main contribution is altering the pipeline to a fc-mer based model, where the traditional read mapping and variant calling stages are replaced by LAVA’s unified genotyping method.

As increases in modern genomic sequencing capabilities have been outpacing even the exponential increases in computing power and storage (Kahn, 2011;Berger *et al.*, 20139), continuing to extract meaningful knowledge from this data deluge requires not only faster computers, but also algorithmic advances. One popular approach has been to accelerate existing tools and algorithms using more sophisticated data structures (e.g. the FM-index (Ferragina *et al.*, 20049), compressive acceleration (Loh*et al.*, 2012;Daniels *et al.*, 2013;Yu *et al.*, 2015a), etc.). However, read mapping is used as a building block for many different downstream applications (Langmead and Salzberg, 2012), so it must be designed to be as general as possible. When the specific downstream application is known beforehand, the read mapping information need not be fully computed giving way to more efficient, ‘alignment-free’ approaches.

While alignment-free sequence comparison has been around for more than two decades (Hide *et al.*, 1994;Jeffrey, 1990;Vinga and Almeida, 2003), their mainstream use in the context of fast processing of large NGS datasets is relatively recent, including tools making use of lightweight alignment (Patro *et al.*, 2015) or pseudo-alignment (Bray *et al.*, 2015) for transcript quantification and metagenomic classification (Wood and Salzberg, 2014;Ounit *et al.*, 2015) as well as fully reference-independent methods that identify differences between wild-type and mutant individuals (NordstrÖm *et al.*, 2013;Peterlongo *et al.*, 2010). These methods differ from traditional genomic pipelines by going from unaligned reads to analysis-ready results without needing to compute nucleotide-level alignments of reads onto a reference sequence. In this paper, we show that a *k*-mer based algorithm that employs similar alignment-free sequence comparison principles, yet allows approximate *k*-mer matches, can accurately genotype an individual for a given set of SNPs.

Due to linkage disequilibrium (LD) between variants that are close in terms of recombination distance, relatively few SNP loci in the human genome are needed for tagging the variants present in an individual (Frazer *et al.*, 2007). As such, a fast algorithm that can compute genotype information of a given set of SNPs, even if it eschews discovery of novel SNPs, is of great relevance in population genomics, impacting ancestral and genome-wide association studies (GWAS). Aptly, fast genotyping methods are most urgently needed in population-level studies, where sequencing data from a large number of individuals need to be processed for analyses.

**Fig. 2.**
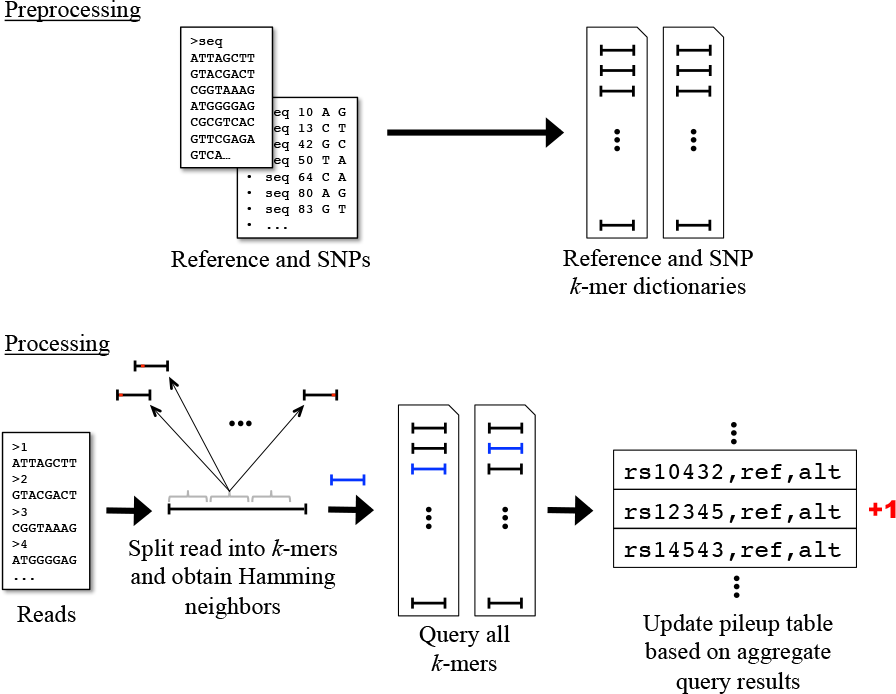
High-level view of LAVA method. We first produce dictionaries of all reference fc-mers and fc-mers containing mutant SNP alleles in the preprocessing stage from the given reference sequence and SNP list. These dictionaries associate a position in the reference with each *k*-mer. The SNP dictionary also contains reference and alternate alleles. The subsequent online processing of the reads entails querying each read’s constituent *k*-mers, in addition to their Hamming neighbors, in these two dictionaries. For each read, the results of these queries are combined in order to predict which SNPs the read overlaps, and we increment either the reference or alternate allele counter in our pile-up table for all such SNPs, depending on which allele the read contains for that SNP. Once all reads have been processed in this way, the final, completed pile-up table is used to call variants.

## 2 Approach

Here we introduce LAVA (Lightweight Assignment of Variant Alleles), which from raw sequencing reads performs something akin to a computational SNP array, calling SNPs as either wild-type or mutant. In particular, given a set of SNPs, LAVA constructs a comprehensive dictionary of mid-size *k*-mers (with *K* = 32) that uniquely identify those SNPs (where possible). Coupled with a second dictionary of all the *k*-mers in the human genome, LAVA is able to quickly determine if a read belongs to a particular SNP as either a wild-type or mutant through a bipartite matching of *k*-mers in the reads to fc-mers in the dictionaries up to Hamming distance 1. The *k*-mers used in LAVA can be seen as the computational analogue of the ASOs used on SNP arrays, allowing us to choose only relevant reads, without doing a full alignment of all reads to a reference genome (Figure 1). By aggregating those relevant reads, LAVA can then call SNPs with a simple probabilitic model using expected read-depth coverage as well as variant frequency priors from dbSNP (Sherry *et al.*, 2001).

LAVA accurately genotypes the vast majority of SNPs in our experiments significantly faster than traditional genotyping through mapping. For a SNP list consisting of a subset of common SNPs from dbSNP, the speedup was 3.4−6.0x. Similarly, using the SNPs from the Affymetrix Genome-Wide Human SNP Array 6.0 as a SNP list, we saw a speedup of 2.2−9.2x. Furthermore, LAVA was able to use as little as ~40GB of RAM for the dbSNP-common SNP list and ~5GB of RAM for the Affymetrix SNP list. At the same time, because LAVA is a computational method that relies on NGS, it does not require the construction of physical SNP chips, and can address many more SNPs than ASOs can feasibly be dotted onto a chip. Moreover, when the set of SNPs is altered, LAVA’s dictionaries can easily be modified in silico as opposed to expensively redesigning the probes used in the SNP array.

**Fig. 3.**
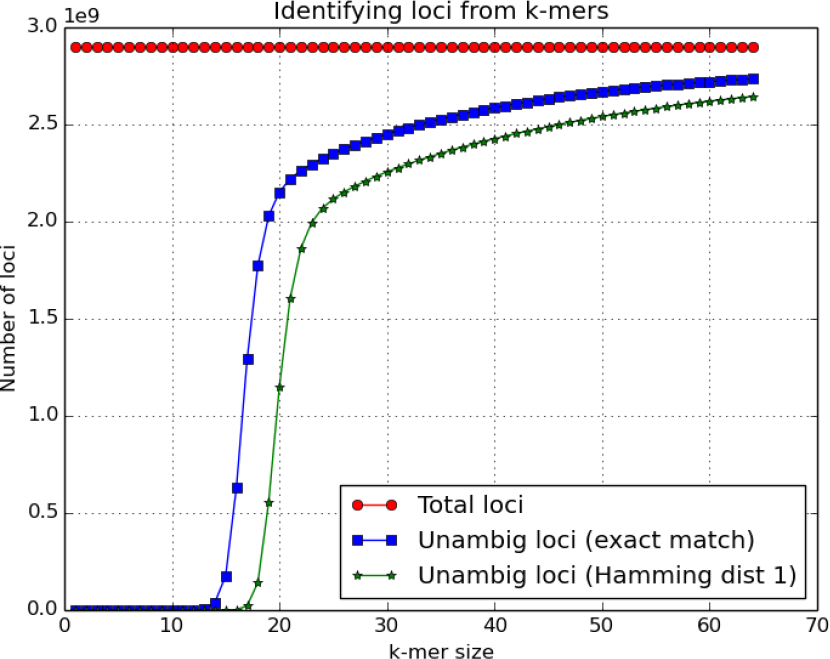
The number of identifiable loci in the human reference from *k*-mers of different lengths. Total number of loci is ~3 × 10^9^ (red circles). Naturally, both the number of unambiguous loci given exact *k*-mer matches (blue squares) and taking into account Hamming neighbors (green stars) increase with *k*. Clearly, in the latter case we need much longer *k*-mer lengths to correct for the presence of errors injecting ambiguity into the loci corresponding to any particular *k*-mer.

Our broader point is that while full mapping and genotyping will be useful for the discovery of novel SNPs and more in depth computational analyses of genomes, it is more costly than necessary for many population genomics studies. LAVA provides a computationally much cheaper solution for genotyping applications on a given set of SNPs.

On our human test dataset (NA12878 from the 1000 Genomes Project (Consortium *et al.*, 2012) and the GATK best practices bundle (DePristo *et al.*, 2011)), LAVA correctly called 93.1% of a subset of common SNPs from dbSNP, and 96.4% of SNPs from Affymetrix Genome-Wide Human SNP Array 6.0. By comparison, the other conventional genotyping pipelines that were tested correctly called 92.6–94.8% of the former and 93.4–95.5% of the latter.

## 3 Methods

LAVA takes as input a reference genome, a list of SNPs, and a set of reads. As its output, it produces predicted genotypes for those SNPs (wild-type, heterozygous, homozygous mutant). A high-level visualization of the method is depicted in Figure 2.

### 3.1 Choice of *k*-mer Length

We choose *k* = 32 for a combination of theoretical and machine architectural considerations. First and foremost, we want our *k*-mers to, as much as possible, uniquely identify the loci of the genome where SNPs of interest are located. This is akin to the problem of the choice of ASO probe sequence in SNP arrays. However, though ASO probes are generally of length between 15 and 21, this choice only works for a small subset of the human genome. Additionally, in order to take into account machine error (which ASO probes need not), we also want to make sure to choose a sufficiently long *k*-mer so that *k*-mers also do not have Hamming neighbors in the genome (loci that are within Hamming distance 1). Figure 3 is an analysis of the human reference genome (version 19), showing how many loci are uniquely identifiable by *k*-mers of different lengths, either with exact matches or when Hamming neighbors are also considered. Only 21.8% of 16-mers are unique in the human reference genome and only 0.000786% of 16-mers have no Hamming neighbors. On the other hand, by choosing *k* = 32, we are able to uniquely locate 85.7% of the human genome with exact 32-mer matches and 79.3% of the human genome have no Hamming neighbors. Of note, although the proportion of uniquely identifiable loci continues to increase with *k*, 32 seems to be past an inflection point and is well-suited for modern 64-bit machine architectures.

On the other hand, too large of a choice for *k* runs into a different set of problems. The most obvious practical problem is that the memory requirements for *k* much larger are prohibitive. Additionally, there is also a more subtle issue caused by sequencing error rates: as *k* grows, the chances of sequencing error being present also grow. By looking to Hamming neighbors of the *k*-mer, we can correct for a single error present in a *k*-mer, but looking to higher Hamming distances requires exponentially more time. Thus, we need to ensure that *k*-mers rarely have more than one machine error. Assuming independence of error locations with an error rate of *p*, the binomial distribution gives that a *k*-mer will have *l* errors with probability

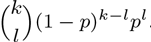

With even a low 1% error rate, *k* = 32 gives ≥ 2 errors at a rate of only 4%, whereas *k* = 64 results in ≥ 2 errors at a rate of 13%. At just a 2% error rate, *k* = 32 gives ≥ 2 errors at a rate of only 13%, whereas *k* = 64 results in ≥ 2 errors at a rate of 36%.

As far as machine architecture is concerned, a 32-mer can be simply encoded as a 64-bit number. Note that *k* = 16 and *k* = 64 also fit nicely in standard machine number sizes, but they do not serve our purposes for the reasons listed above. Thus, we have chosen *k* = 32, which fits all of the criteria above relatively well.

### 3.2 Preprocessing of Reference Sequence and SNP List

We begin by preprocessing the given reference sequence by considering every *k*-length substring (“*k*-mer”) appearing within it. Our goal is to create a dictionary D_ref_ that maps each *k*-mer to the index in the reference sequence at which the *k*-mer appears. If a given *k*-mer appears more than once in the reference sequence, we can treat it specially and store multiple positions for it, up to some limit.

Using a standard binary encoding of reads, let ξ be the bijection mapping each 32-mer to its 64-bit unsigned integer encoding. Simply treating *D*_ref_ as a list of (*k*-mer, position) tuples does not allow for efficient querying. To decrease cache misses, we use a static hash table implementation as follows (Yu *et al.*, 2015b): we first sort *D*_ref_ by the numerical values of the encoded *k*-mers, and then we make use of a secondary hash table *J*_ref_ that maps each 32-bit unsigned integer *u* to the first location in *D*_ref_ at which there is a 32-mer whose encoding’s upper 32 bits is u. Because *D*_ref_ is sorted by the numerical values of the encoded 32-mers, 32-mers whose encodings have the same upper 32 bits will be grouped together in a sorted bucket, decreasing cache misses when searching all Hamming neighbors of a *k*-mer and allowing us to implicitly encode the upper 32 bits of *k*-mers by bucket, improving memory efficiency. To query *D*_ref_ with some 32-mer *K*, we simply binary search in *D*_ref_ between the indices *J*_ref_[└ξ(*K*)/2^32^┘] (inclusive) and *J*_ref_[└ξ(*K*)/2^32^┘ + 1] (exclusive) for *k*. A simplified visualization of the querying process is given in Supplementary Materials.

The known SNP list is also preprocessed analogously: instead of considering all *k*-mers in the reference, we take only those that overlap some SNP with the reference allele replaced by the alternate. Each element of the SNP list consists of a position in the reference at which the SNP is located, a reference allele, an alternate allele and population frequency priors of the reference and alternate alleles. The SNP dictionary D_SNP_ (as well as a secondary hash table *J*_SNP_) thus contains *k*-mers (*k* = 32 again) with the mutant allele. In addition to positions, we store SNP information in *D*_SNP_ for each *k*-mer.

Though these structures are constructed during preprocessing, they still must be loaded into **RAM**. We give exact total numbers in Table 1, but space complexity of the static data structures is at most linear in size of the genome (because each locus in the genome and each SNP contributes one *k*-mer and each *k*-mer stores no more than constant information for numbers of coinciding loci). Alternately, if only relevant *k*mers are stored, one can bring the space complexity down to linear in the number of SNPs of interest (see LAVA Lite in Table 1). Additionally, the constant overhead of the hash table buckets can be tuned to be on the order of the number of *k*-mers.

### 3.3 Online Processing of Reads

To obtain variant calls from reads, we first create a “pileup table” *P*, where we store for each SNP, reference and alternate allele counts (respectively α and β), which are updated incrementally as we process the reads. *P* can be thought of as a dictionary mapping indices to SNPs and these two counts. This pile-up table can be implemented to use space linear in either, trivially, the genome length, or using a hash table, the number of SNPs, so the space-complexity of LAVA is determined by the static structures given in the previous section.

Each individual read *Q* can be thought of as a length-*m* sequence of A,C,G,T (N’s are handled separately as we discuss below). The processing of a read begins with splitting it into non-overlapping, contiguous *k*-mers. If there is a segment at the end of the read that is not covered by one of these *k*-mers, we can optionally either omit that segment or choose a final overlapping *k*-mer that covers it.

When the sequencer cannot call a base and emits an N, there are several possible courses of action. The simplest one, and the one we use in the results presented in this paper, is to discard any read that contains an N. Alternately, as the LAVA framework is flexible with respect to read length, another practical solution is to trim a read if N’s are clustered near the end of the read. Lastly, for sporadic N’s in the middle of the read, another option is to expand our alphabet to include N for use in Hamming distance computations, but not allow it to play a role in SNP calling.

Now, let*𝒩*(*K*) be the “Hamming neighborhood” of some *k*-mer *k* for Hamming distance 1. Notice that *K* ∈ *𝒩*(*K*) and that *|𝒩*(*K*)| = 3*k* + 1. Our immediate goal in processing each read is to determine which SNP(s), if any, the read corresponds to. We can then update the reference and alternate allele counts in *P* for such SNPs appropriately. We identify these SNPs by querying *D*_ref_ and *D*_SNP_ with all *k*-mers in *𝒩*(*K*) for each *k*-mer sampled from the read. For each read, we identify the locus to which the most *k*-mers in that read can be concordantly matched. If there are multiple such loci, or if there is no locus supported by two or more *k*-mers, we do not update *P*. We apply this entire procedure to each read and to its reverse complement, resulting in a completed pile-up table *P\**.

### 3.4 SNP Calling

The final stage of LAVA is to utilize the pile-up table *P\** to call the donor’s genotype for each SNP locus. Specifically, we assign a label of either “homozygous reference” (*G*_0_), “heterozygous” (*G*_1_) or “homozygous alternate” (*G*_2_) to each position in *P\** that is covered by at least one read (i.e. α + β > 0). Since we have the reference and alternate allele frequencies (*p* and *q* respectively) in *P\**, we can estimate the posterior probability of each genotype as:

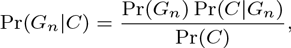

where *C* is the event that we observe reference and alternate allele counts of α and β, respectively. Furthermore, by the law of total probability, we have

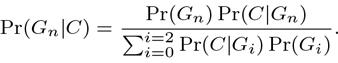

By Hardy-Weinberg, we know that

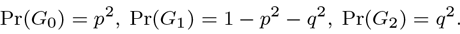

We take each Pr(*C|G_n_*) to be binomially distributed. For *G_0_* and *G_2_*, we assume that we observe an incorrect allele (i.e. alternate allele given *G_0_* or reference allele given *G_2_*) with some probability ε. For *G_1_*, on the other hand, we assume that we have an equal chance of seeing both the reference and alternate alleles. Hence, we have

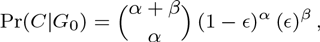

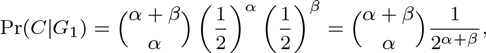

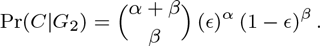

With this observation, we are able to compute each Pr(*G_n_|C*). We take whichever *G_n_* produces the largest Pr(*G_n_|C*) to be our predicted genotype for the given SNP. As our confidence metric for this assessment, we take the Pr(*G_n_|C*) value scaled by a quantity that depends on the total coverage, α+ β In this way, we penalize SNPs that have abnormally high coverages as well as those that have lower coverages than expected. This scaling term is simply the probability mass function of a Poisson distribution with mean equal to the average coverage λ of the reads:

**Table 1.**
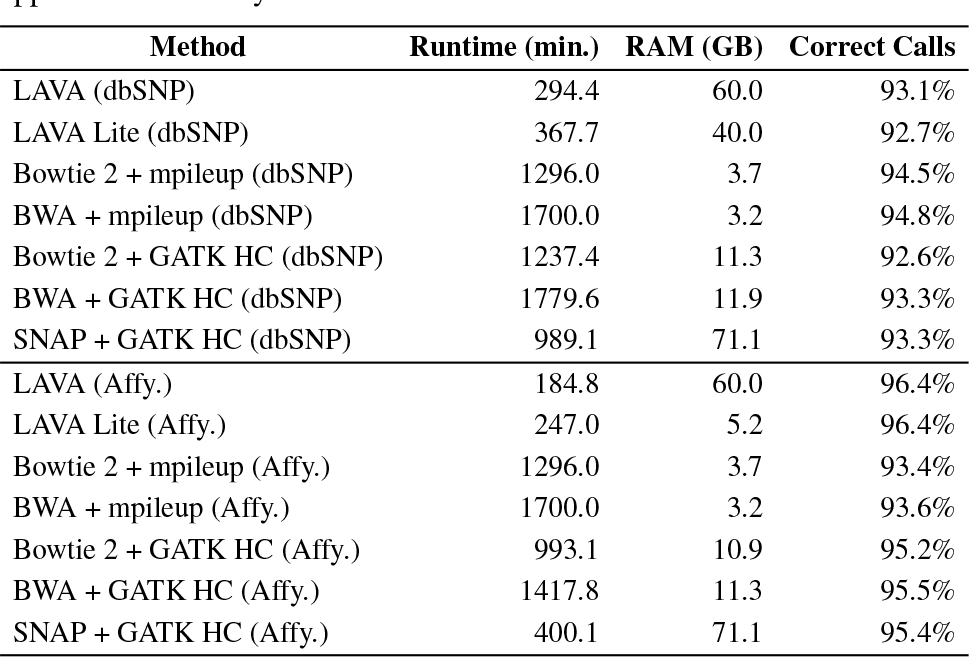
Table of overall running times, peak memory usages, and accuracies for LAVA and for several other genotyping pipelines. Results are shown for two SNP lists: a subset of common SNPs from dbSNP and the SNPs from the Affymetrix Human SNP Array 6.0. Each tool was allocated a single thread on an Intel Xeon E5-2650 x86_64 CPU @ 2.30GHz. Note that LAVA Lite attains a lower peak memory usage by removing *k*-mers from the reference dictionary that are not within a read length of any SNP, condensing the pile-up table, and (for the Affymetrix SNP list) using 24-bit keys in the reference dictionary as opposed to 32-bit keys.

For greater accuracy, we computed the average α + β from*P\** to measure the actual observed depth coverage, and took λ to be this value. We report the product of these two probabilities as our genotyping confidence values.

**Fig. 4.**
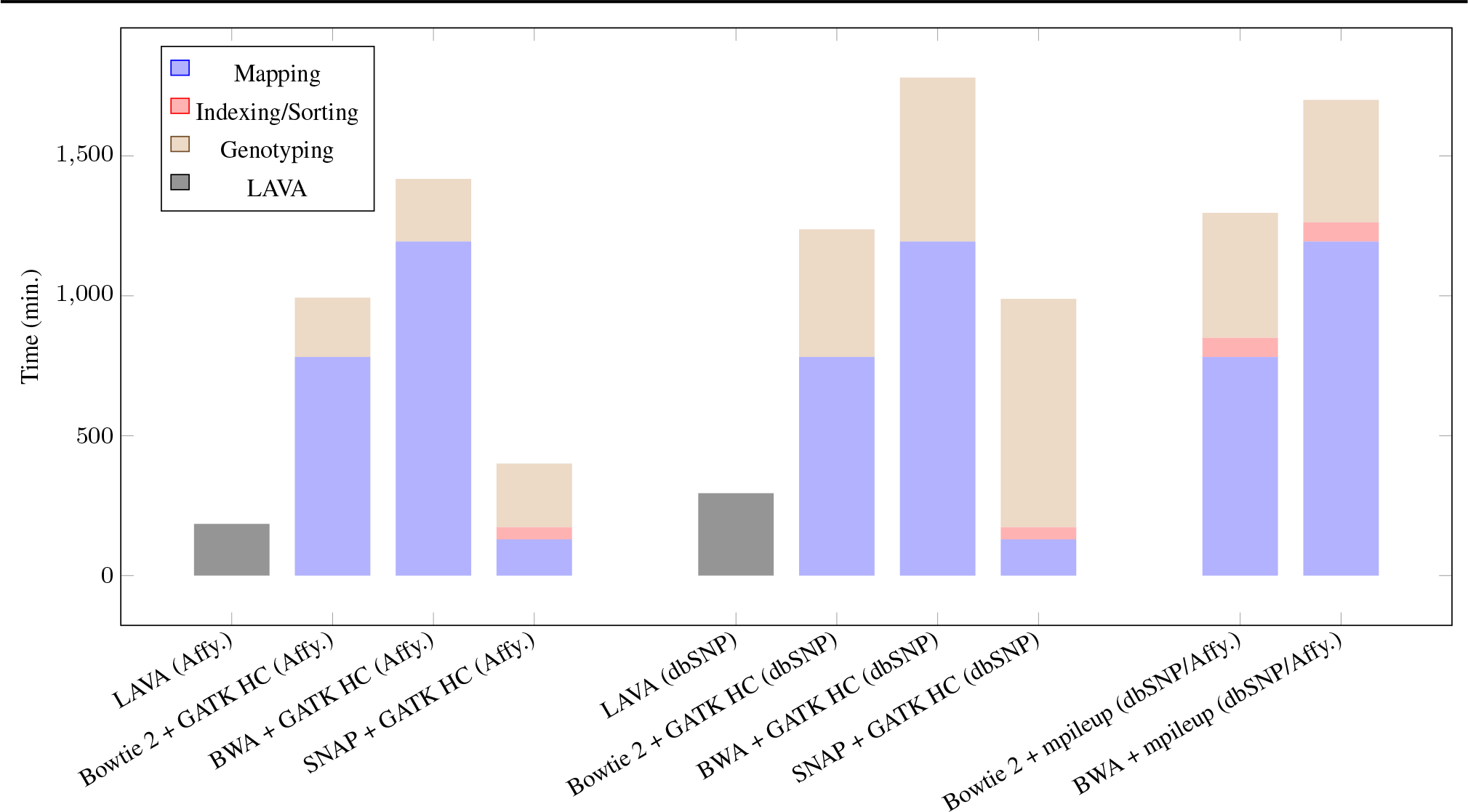
Visualization of LAVA runtimes as compared to other genotyping pipelines. Note that “Indexing/Sorting” refers to intermediate operations performed on the output of the mapping stage prior to genotyping. The actual numerical values of the various times are given in Supplementary Materials.

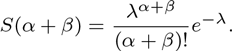

### 3.5 Parallelization

Parallelizing LAVA is straightforward, as different reads can be processed independently by different threads. For comparison, a parallelized version of LAVA (using a subset of common SNPs from dbSNP as the SNP list) running with 4 threads (using shared memory) achieved a 3.3x speedup while a pipeline consisting of Bowtie2 for read mapping and GATK’s HaplotypeCaller for genotyping had a 2.9x speedup.

## 4 Results

*Datasets.* As our reference sequence, we used GRCh37/hg19. We used NA12878 reads from Phase 1 of the 1000 Genomes Project (Consortium *et al.*, 2012) and a high-quality trio-validated genotype annotation as our gold standard (DePristo *et al.*, 2011). Then we performed the experiment for two different SNP lists: all common SNPs from dbSNP (Sherry *et al.*, 2001) and SNPs from the Affymetrix Genome-Wide Human SNP Array 6.0. We compared our accuracy with the most popular genotyping pipelines, consisting of various combinations of Bowtie 2, BWA or SNAP for read mapping and Samtools mpileup or GATK’s HaplotypeCaller (henceforth also referred to as “GATK HC”) for variant calling. Additionally, as the HaplotypeCaller allows the user to specify a set of alleles at which to genotype, GATK HC was also run specifically for the Affymetrix SNP list with this setting enabled. Note that each SNP list is filtered so that it contains only bi-allelic, single-nucleotide SNPs with consistent alleles and allele frequency data.

*Parameters.* For the Poisson distribution in our model, we set λ = 7.1, which was the average coverage in our final pile-up table. Our assumed error rate was ε = 0.01, which is in line with known NGS base call error rates. For *k*-mers with multiple mappings in the two dictionaries, we stored up to 9 additional entries. Any reads containing an N base (~0.5% of all reads) were discarded. Also, segments at the end of a read not evenly covered by a *k*-mer were also discarded; for our read length of 101 bp, this corresponded to discarding the last 5 bases in each read, which were also the lowest quality regions of the read.

*Benchmarking experiments*. Depending on mode, LAVA used from as little as 5.2 GB to up to 60 GB of peak memory while running for our dataset. By contrast, BWA + mpileup used about 3.2 GB at its peak, Bowtie 2 + GATK HC used 11.3 GB, and SNAP + GATK HC used 71.1 GB. In Table 1, we present the timings and peak memory usages for all of our experiments, both for LAVA and for the other genotyping pipelines. LAVA allows for trading off memory usage for runtime. Of note, even the lowest memory mode achieves impressive speed gains (Table 1 and Figure 4).

*Dictionary generation*. Generating the reference and SNP dictionaries took a combined time of about 28 minutes on our machine (described in the caption of Table 1), and used about 74 gigabytes of memory at its peak (these benchmarks are essentially independent of the SNP list since generating the reference dictionary is predominantly the time-and memory-consuming step). Note that this is a preprocessing stage and that the same reference and SNP dictionaries can be used repeatedly for the same reference genome and SNP list, respectively.

Overall, compared to the other genotyping pipelines, LAVA proved to be anywhere from 3.4 to 6 times faster for the dbSNP-common SNP list and anywhere from 2.2 to 9.2 times faster for the Affymetrix SNP list. Figure 5 shows the accuracy of LAVA and that of the other pipelines for the two different SNP lists.

**Fig. 5.**
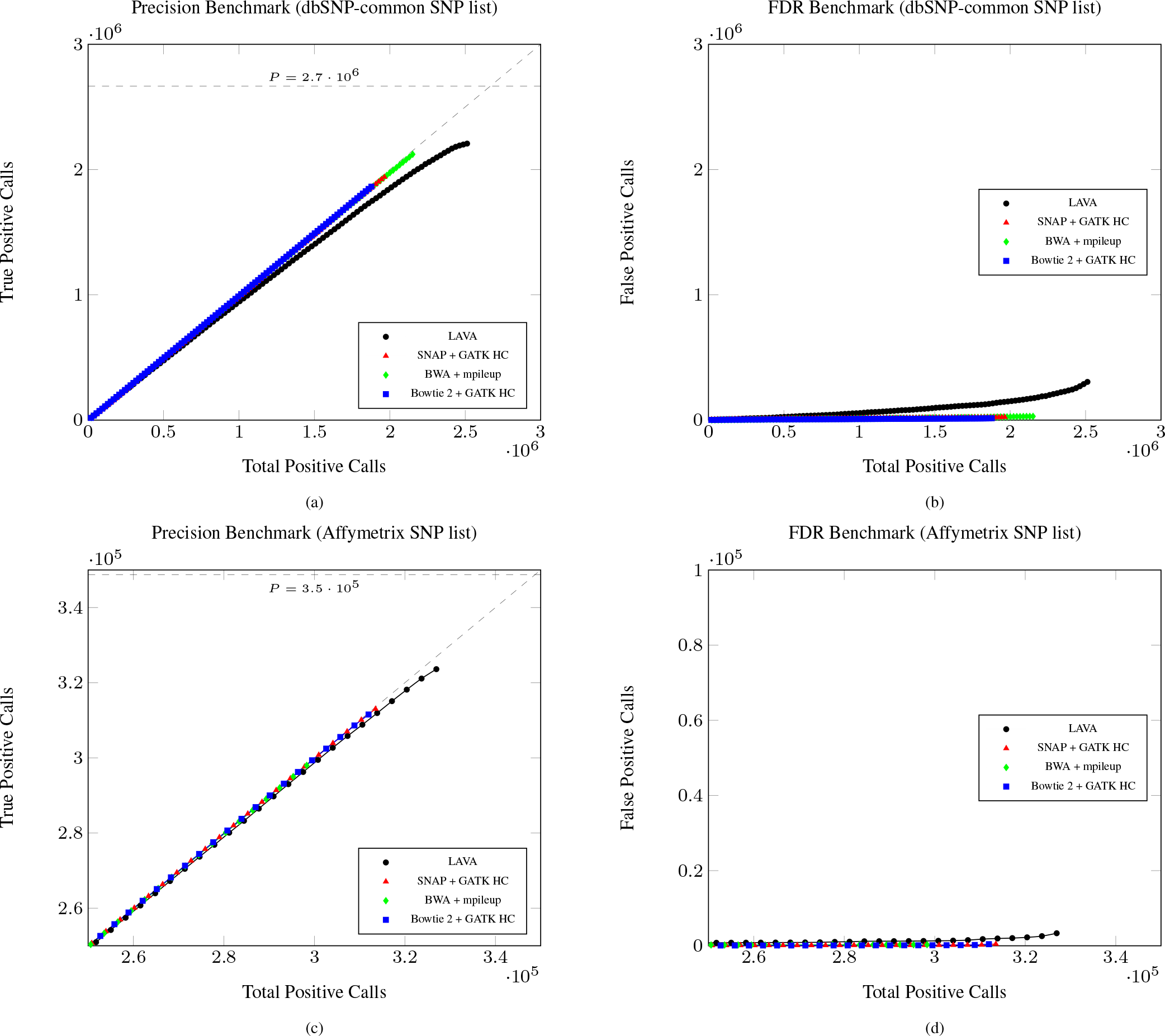
Accuracy plots for the dbSNP-common SNP list as well as for the Affymetrix SNP Array 6.0 list, showing true positives (a,c) and false positives (b,d) as a function of total positive calls, both for LAVA and for several other genotyping pipelines. P denotes the total number of positives present in the dataset. Note that the positive set contains the heterozygous and homozygous mutant SNPs (*G*_1_ and *G*_2_) and the negative set contains the homozygous wild-type SNPs (*G*_0_).

Because a small subset of SNPs cannot be uniquely identified by any of their overlapping 32-mers, LAVA only attempted to call 94.5% of dbSNP-common and 98.5% of the Affymetrix SNPs. LAVA correctly called 93.1% (98.5% of attempted calls) of the dbSNP-common list, and 96.4% (97.9% of attempted calls) of the Affymetrix SNP Array 6.0 list. The Bowtie 2 + mpileup pipeline, by comparison, correctly called 94.5% and 93.4% respectively. Furthermore, BWA + mpileup called 94.8% and 93.6% while Bowtie 2 + GATK HC called 92.6% and 95.2%. Hence, despite its speed, LAVA maintained a high accuracy for the dbSNP-common SNP list and actually attained the *highest* accuracy for the Affymetrix SNP list.

Though LAVA performs genotyping from raw reads in a single unified algorithm, rather than separating out mapping and variant calling, its fast speed compares favorably against the individual components of existing pipelines. As such, even when compared to variant calling from a precomputed BAM file with no cost spent on mapping, LAVA can provide significant speed improvements.

## 5 Discussion

LAVA applies the idea of mid-size *k*-mer based lightweight algorithms to the problem of genotyping and in doing so achieves great improvements in speed. By replacing full read mapping and variant calling from the sequence analysis pipeline with LAVA, we improve the speed of the process and also unify it considerably in a framework that performs an approximate bipartite matching between *k*-mers in the reference and the read datasets.

While in this study we have focused LAVA on SNPs, DNA microarrays are applicable to other variants such as insertions, deletions, and CNVs (copy number variations). Because physical ASOs (with 15 ≤ *k* ≤ 21) can assay for these variants, we expect that even accounting for sequencing error, the 32-mers LAVA uses can also address these variants. For small indels, the reference and alternate *k*-mers are exactly analogous to the ones used for SNPs, requiring only that we find nearly unique *k*-mers for each variant. With a length-*n* insertion, for instance, we would take the *k* (*k* – *n*)-mers surrounding the start of the insertion and place the insertion sequence into each of them at the correct position to produce *k k*-mers representing the insertion. Similarly, for a length-*n* deletion, we would take the *k* (*k* + *n*)-mers surrounding the start of the deletion and remove the deleted sequence from each to produce *k k*-mers representing the deletion. Copy number variation information is already present to some extent in the pile-up table, and we need only update the Bayesian model to include more possible classifications; it is likely that CNVs will require higher average coverage to accurately call. Or, alternately, known structural duplications may be accessible through *k*-mers covering the boundary between a duplicated region and its neighboring regions.

Additionally, in the interest of speed, LAVA uses a more streamlined probabilistic base calling algorithm. However, once LAVA performs its *k*-mer matching step, any number of different probabilistic models could be applied for base calling. Further augmentations to the LAVA framework could include more complicated models—such as those used in tools like GATK HaplotypeCaller and Samtools mpileup—and better priors for the Bayesian model, such as linkage disequilibrium information. While these other models have some advantages in accuracy, we deliberately chose to make the trade-off of gaining additional speed through our streamlined probabilistic model, while still retaining reasonably high accuracy.

Of note, LAVA demonstrates the highest accuracy amongst all pipelines we tested for the Affymetrix dataset. This is because LAVA’s dictionaries effectively simulate the behavior of ASO probes, so SNPs for which SNP arrays are effective are also particularly amenable to LAVA’s variant calling.

## 6 Conclusion and Future Work

While we demonstrate LAVA’s capabilities for efficiently processing DNA sequencing datasets for known SNP loci, if common splice junction coordinates in the population are known, which is available from studies such as GENCODE (Derrien *et al.*, 2012) and REFSEQ (Pruitt *et al.*, 2007), LAVA can also be augmented to perform bipartite matching of *k*-mers in the RNA-seq reads and the transcriptome for identifying SNP/indel variants as well as estimate differential allelic expression of heterozygous loci.

Furthermore, although paired-end reads were used in our benchmarking experiments, LAVA does not actually distinguish between paired-and single-end, unlike the read mappers we compared against. Future work on LAVA entails incorporating a paired-end mode to further improve accuracy.

To conclude, as NGS read databases grow, the lightweight nature of LAVA will enable much faster targeted analyses than can be performed with standard NGS genotyping pipelines. These methods will prove invaluable as sequencing becomes the assay of choice for population genomics studies.

## Acknowledgements

We thank Noah Daniels, Hoon Cho, Sean Simmons, and David Rolnick for helpful discussions and comments.

